# A new method for long-read sequencing of animal mitochondrial genomes: application to the identification of equine mitochondrial DNA variants

**DOI:** 10.1101/2019.12.20.884486

**Authors:** Sophie Dhorne-Pollet, Eric Barrey, Nicolas Pollet

## Abstract

**Background:** We present here an approach to sequence whole mitochondrial genomes using nanopore long-read sequencing. Our method relies on the selective elimination of nuclear DNA using an exonuclease treatment and on the amplification of circular mitochondrial DNA using a multiple displacement amplification step.

**Results:** We optimized each preparative step to obtain a 100 million-fold enrichment of horse mitochondrial DNA relative to nuclear DNA. We sequenced these amplified mitochondrial DNA using nanopore sequencing technology and obtained mitochondrial DNA reads that represented up to half of the sequencing output. The sequence reads were 2.3 kb of mean length and provided an even coverage of the mitochondrial genome. Long-reads spanning half or more of the whole mtDNA provided a coverage that varied between 118X and 488X. Finally, we identified SNPs with a precision of 98.1%; recall of 85.2% and a F1-score of 0.912.

**Conclusions:** Our analyses show that our method to amplify mtDNA and to sequence it using the nanopore technology is usable for mitochondrial DNA variant analysis. With minor modifications, this approach could easily be applied to other large circular DNA molecules.

## Background

Mitochondria have become crucial organelles of eukaryote cells since an ancestral bacterial endosymbiosis event gave rise to the eukaryote branch of the tree of life, about 1.5 billion years ago (1,2). The discovery that mitochondria contains DNA in the early 1960s attracted the attention of many scientists and more recently the word of mitogenomics has been coined to unite this field of study (3). Today, thousands of mitochondrial genomes (mtDNA) from many species have been sequenced, and ongoing studies require robust, rapid and cheap method to decipher novel genomes and identify genetic variants (4). In human alone, more than 48,882 full length mtDNA sequences are known, but much less sequences and variants are available for other mammals (5). Numerous and diverse applications take advantage of mtDNA sequencing, such as diagnostic of mitochondrial diseases in human and animals, breeding selection, forensic studies, biodiversity surveys, conservation genetics and evolutionary analysis.

Animal mitochondrial genomes are usually encoded by a single circular DNA of about 10 to 20 kbp, but weird cases are known such as linear or very large molecules (6). The mitochondrial genomes are characterized by a maternal transmission, a lack of recombination and higher frequency of mutations than in the nuclear genome (7). In vertebrates, the ratio of mtDNA over nuclear DNA mutation rate is usually above 20 (8). This means that mtDNA polymorphisms may occur relatively often in animal populations with the potential for associated disorders, and this is why sequencing these organelle genomes is an important component of contemporaneous animal genetics. Knowing the whole mtDNA sequence enables the identification of all possible genetic variants: single-nucleotide polymorphisms (SNP), insertions, deletions, duplications, inversions, rearrangements. Although mitochondrial variants are typically maternally inherited, they can be sporadic. Such *de novo* variants are difficult to assess due to the co-existence of several mtDNA population in a cell (heteroplasmy) that may be below an assay’s detection level, or different between tissues of the same organism (9). The presence of DNA segments transferred from the mitochondria to the nuclear DNA, the so-called Nuclear Mitochondrial DNA Sequences (NUMTs), can seriously complicate mtDNA analysis in a variety of animals from bees to humans (10–12). Such NUMTs can span several kilobases, can be nearly identical to mtDNA sequence and can exhibit variation between individuals of the same species (13).

The current methods for mtDNA sequencing rely on standard sequencing methods such as Sanger sequencing or massively parallel sequencing (4). The choice of the starting DNA material is of importance since the relative mass amount of mtDNA is very low in comparison to nuclear DNA. Typical DNA preparations comprise approximately 0.1% mtDNA, even though there are hundreds to tens of thousands copies of mtDNA per cell (14). One strategy relies on the amplification of mtDNA using regular or long-range PCR. This process is associated with an unpredictable risk of PCR errors, it can introduce a sequence bias, and it can be tricky because it depends on primers, on PCR efficiency, and finally the whole procedure can be time consuming and costly. In addition, the reliance on PCR primers derived from a reference sequence can drive the failure of some amplifications because they may suffer from a lack of specificity to mtDNA, and they may lead to the production of artefactual amplicons, including some derived from NUMTs (15–17). The other strategy rely on whole-genome shotgun using massively parallel sequencing approaches, an efficient approach for mitogenomics because mtDNA reads are more prevalent than nuclear DNA reads, but it is not bullet-proof and short read mapping can also be confounded by NUMTs (18,19). These approaches involve significant resources for bench work and efforts in sequence handling, they are cost-effective only when high number of samples are pooled and therefore they do not scale down easily (18). The principal caveat of the current mtDNA sequencing methods is that raw sequences enable to read only a tiny portion of the DNA at a time, preventing definitive evidence for long-range phasing or structural variant identification.

Long-read sequencing using Oxford Nanopore or PacBio technologies provide the possibility to capture in a single read the majority or the whole sequence of mtDNA molecules. Therefore, it can provide definitive evidence that a given mitogenome exists in a single cell or in a sample. So far, a few studies harnessed the Nanopore technology for sequencing mtDNA. Istace and colleagues assembled 21 yeast mitogenomes in a study where they evaluated the performances of nanopore-only assemblers. They used only 2D reads for mitogenome assembly, a chemistry that provided better sequence quality but that has been abandoned (20). Ranjard and collaborators assembled the green-lipped mussel mitogenome (*Perna canaliculus* Gmelin 1791) using a whole-genome shotgun approach with long-reads to obtain a draft assembly that was polished using short-reads from a RNAseq experiment; but they do not give any details on the long-read sequencing output (21). In another study, the european lobster (*Homarus gammarus*) mitogenome was assembled separately with both Illumina short-reads and Nanopore long-reads (22). The report by Gan and collaborators is instructive since they could not obtain a large mtDNA contig using de novo or single gene bait-based assemblies of six million Illumina paired-end reads totalling 1.5 Gbp. However, the 20 kbp lobster mitogenome could be assembled from 40,687 long-reads totalling 100 Mbp. The same authors reported another mitogenome assembled similarly, with only nine long-reads (23). These reports testify the usefulness of the Nanopore technology to sequence mitochondrial genomes, but the corresponding protocols are not tailored to identify mtDNA variants on a large-scale.

In horses, many mitochondrial functions have been shown to play important roles for muscular exercise and disorders (24–27). Yet, the study of mtDNA variation is lagging behind even though it has been instrumental to study ancient horse’s DNA and unravel the domestication events of the wild horse (28–30). As in other mammals, the abundance and polymorphism of NUMTs is a characteristic of the horse genome and this has negative impact on the reliability of mtDNA variant identification (12,31,32).

The goal of our study was to trial a new method for mtDNA resequencing using the long-read technology provided by Oxford Nanopore Technologies (ONT) with the goal of identifying mtDNA variants. We developed an optimised method for mtDNA sequencing starting from total genomic DNA obtained using routine extraction methods. We implemented a protocol based on a nuclear DNA removal by nuclease treatment followed by mtDNA enrichment using multiple displacement amplification (MDA) and sequencing using the 1D chemistry on the MinION platform of ONT. We anticipate that this method will ease the identification of mtDNA variation.

## Results

### Enrichment of circular mtDNA by exonuclease V treatment

We started our strategy by enriching the circular forms of animal mtDNA in genomic DNA extracts. Since traditional biochemical purification of mitochondria does not scale up easily and requires a large amount of fresh tissue, we selected an alternative method. We chose an exonuclease treatment to deplete linear DNA fragments and thereby improve the proportion of mtDNA versus nuclear DNA, as previously described (33). A frequently used enzyme to degrade linear DNA molecules is the exonuclease V that has different nuclease activities, comprising an ATP-dependent double-stranded and bi-directional exonuclease activity. Since the amount of nuDNA vastly exceeds that of mtDNA, we tried four conditions of nuclear DNA removal and evaluated their efficiency using qPCR (Figure 1A). The effect of exonuclease V treatments impacted dramatically the quantity of nuDNA, since it diminished by several thousand fold relative to mtDNA. mtDNA quantity diminished slightly after the various exonuclease treatment to only about half of its amount in the starting DNA extract. We observed that increasing the incubation time had an effect while the amount of exonuclease V seemed not to be a limiting factor. We found that the best condition was two hours of incubation with 10 Units of Exonuclease V, since it resulted in a ratio of mtDNA over nuDNA of 3755+/−0.7 (Figure 1A).

**Figure 1.**
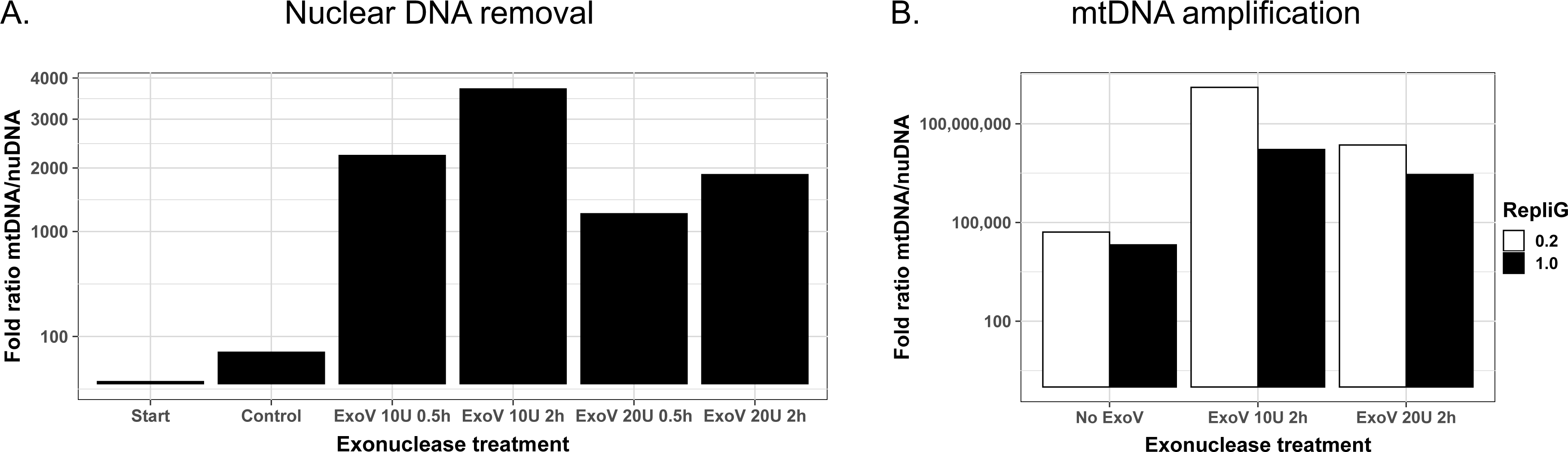
Quantification of mtDNA enrichment by qPCR. A. Quantification after nuclear DNA removal using exonuclease V (ExoV) as indicated. B. Quantification after mtDNA amplification using MDA. The results are given for DNA untreated by exoV (No ExoV), or after two conditions of ExoV digestion as indicated. REPLI-g was performed using primers at a concentration of 0.2 µM or 1.0 µM.

Such an enrichment was of interest, but it came at the cost of a limited amount of DNA which is inadequate for sequencing without an amplification step. We decided to amplify the mtDNA using a Whole Mitochondrial Genome Amplification (WMGA) method based on multiple displacement amplification (34). We optimized primer design and concentration in the MDA reaction to avoid template-independent amplification, a technical bias usually observed with MDA. To evaluate the impact of the Exonuclease V treatment and the amplification efficiency of WMGA, we estimated the relative amounts of mtDNA versus nuDNA with or without an exonuclease V treatment, and using two primer concentrations. We present our results on the quantification of the amplification efficiency in Figure 1B. We observed a nearly 1,000 fold better amplification using WMGA on DNA treated with exonuclease V versus the starting undigested DNA (Figure 1B), with an overall amplification of more than 100 million fold in the best condition using 0.2 µM of each primer. We obtained on the order of ten micrograms of DNA after WMGA, with an average fragment size of about 3 kbp that we deemed usable for long-read sequencing (SI_Figure 1).

### Long-read sequencing

We performed a first sequencing run on nine DNA samples obtained from three horses (H25, H26 and H27) after WMGA on DNA treated or not by exonuclease V and amplified using the two primer concentration conditions mentioned previously. We obtained a total of 634,062 reads with a mean length of 2,370 nt and a mean quality of 10.2. After demultiplexing, we obtained 562,981 reads (88.8 %) evenly distributed among the nine barcodes; with between 45,641 reads and up to 67,448 reads (Table 1 and SI Table 1). We did not observe gross differences of read size or quality between the nine barcoded libraries (Figure 2, Table 1). We mapped these reads to the reference horse genome and counted the reads mapping to the mitochondrial genome and those mapping to the nuclear genome (Figure 2B). The lowest proportion of mtDNA reads, about 1%, was from the DNA samples treated with exonuclease and amplified using the highest primer concentration (exo B, Figure 2B). In the DNA samples without exonuclease treatment but amplified using the lowest primer concentration (exo -, Figure 2), we observed around 5% of mtDNA reads. The proportion of mtDNA reads was up to ten times higher in the DNA samples treated with exonuclease and amplified using the lowest primer concentration (exo A, Figure 2), with values ranging from 17,569 to 40,888 reads, i.e. 30 to 59% of the total sequences from a given sample. Upon inspection of the location of reads mapping to the nuclear genome, we found that about 20% of them were mapped to NUMTs. The effect of the exonuclease treatment and the amplification on the mapping of reads to mtDNA versus nuDNA was significant (Chi2=12552.68, p<0.001). The distribution of read size and quality was similar whether they mapped to the mitochondrial or to the nuclear genome (SI Figure 2). In conclusion, the protocol we developed to amplify mtDNA led to a significant increase in the proportion of long sequences derived from the horse mitochondrial genome.

**Table 1:** Sequencing statistics

| Sample Barcode | H25 exo - Barcode01 | H26 exo - Barcode02 | H27 exo - Barcode03 | H25 exo A Barcode04 | H26 exo A Barcode05 | H27 exo A Barcode06 | H25 exo B Barcode07 | H26 exo B Barcode08 | H27 exo B Barcode09 | Unclassified | Total |
| --- | --- | --- | --- | --- | --- | --- | --- | --- | --- | --- | --- |
| Mean read length (nt) | 2 209 | 2 248 | 2 004 | 2 733 | 2 526 | 2 553 | 1 821 | 3 241 | 2 869 | 2 145 | 2 379 |
| Median read length (nt) | 1 706 | 1 763 | 1 579 | 1 918 | 1 815 | 1 865 | 1 440 | 2 336 | 2 172 | 1 545 | 1 741 |
| Read length N50 (nt) | 2 786 | 2 807 | 2 462 | 3 867 | 3 409 | 3 414 | 2 172 | 4 635 | 3 873 | 2 909 | 3 137 |
| Number of reads | 60 619 | 67 448 | 60 729 | 59 028 | 69 453 | 55 892 | 88 175 | 45 641 | 55 996 | 71 073 | 634 054 |
| % of Total reads | 10% | 11% | 10% | 9% | 11% | 9% | 14% | 7% | 9% | 11% | 100% |
| Total bases (nt) | 133 906 674 | 151 599 799 | 121 723 567 | 161 303 032 | 175 444 343 | 142 699 042 | 160 526 846 | 147 900 066 | 160 661 882 | 152 461 373 | 1 508 226 624 |
| % of Total bases | 9% | 10% | 8% | 11% | 12% | 9% | 11% | 10% | 11% | 10% | 100% |
| Mean read quality | 11.5 | 11.5 | 11.5 | 10.9 | 10.9 | 10.9 | 11.3 | 11.4 | 11.5 | 6.8 | 10.8 |
| Median read quality | 11.7 | 11.8 | 11.7 | 11.1 | 11.2 | 11.1 | 11.6 | 11.7 | 11.8 | 5.8 | 11.3 |

**Figure 2:**
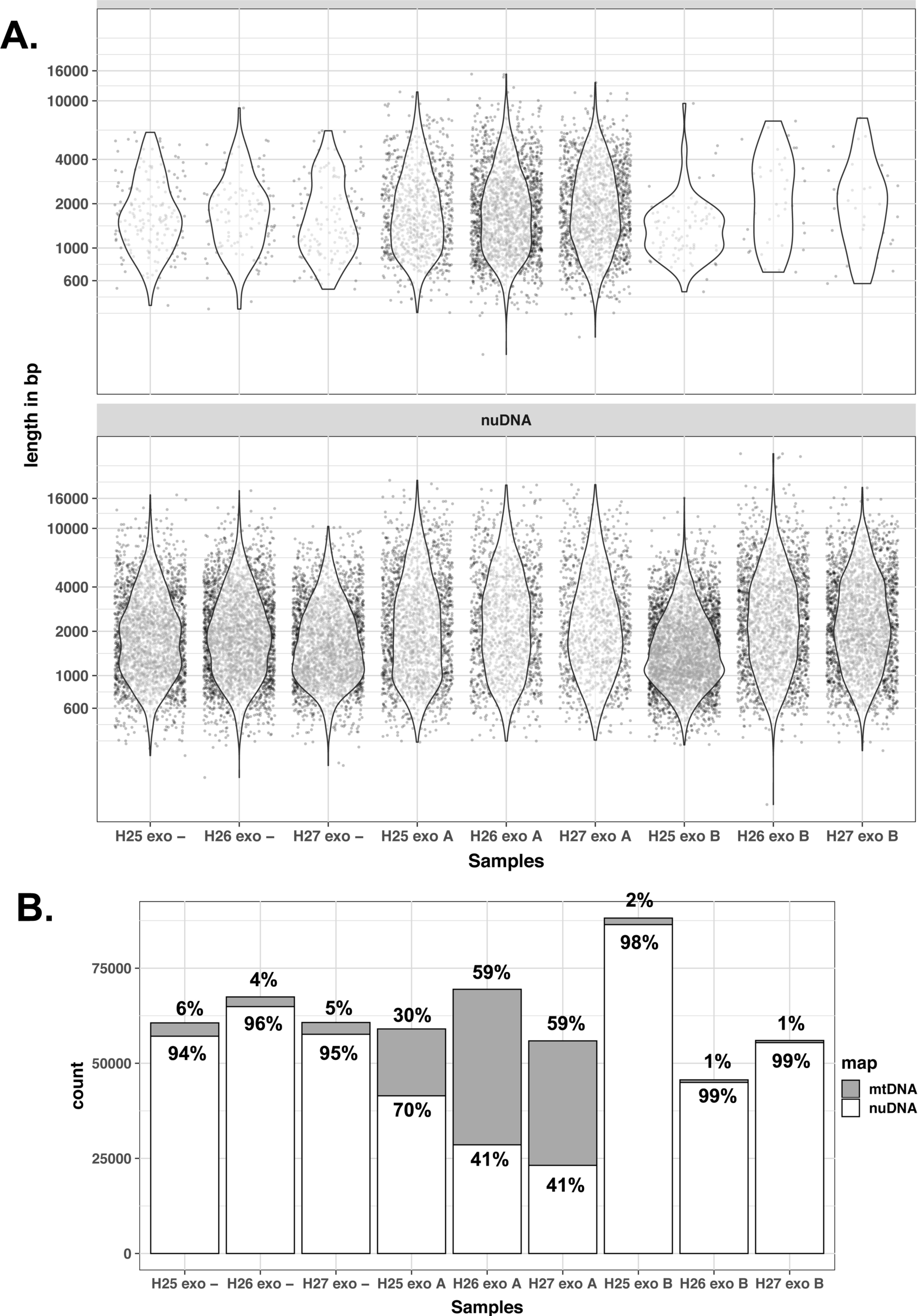
Read length distribution and proportions of sequences mapping to the mitochondrial or nuclear genomes. A. Distribution of reads length mapping to mtDNA (top) or nuDNA (bottom). For clarity, values are shown for a randomly selected subset representing 0.005% of reads B. Numbers and proportions of sequences mapping to mtDNA or nuDNA. The number of mapped sequences is shown on the y axis. exo - : no Exonuclease V treatment ; exo A : Exonuclease V 10 U 2h + REPLI-g 0.2 µM primers ; exo B : Exonuclease V 10 U 2h + REPLI-g 1.0 µM primers.

### Overall coverage of the mitochondrial genome

We then looked at the coverage of the mitochondrial genome at the base level (Figure 3). Whatever the treatment, we found that the coverage of the mitochondrial genome was complete and this pattern was reproducible between the samples. We could observe the effect of some primers on the coverage plot that appeared as bulges, for example between positions 13946 and 14748. For samples untreated by exonuclease in the S1 sequencing run, the minimum coverage was 66 reads at a given mitochondrial base (SI Table 1). In the same run and for the samples treated with exo A the minimum coverage was 526 reads. The variation between minimal and maximal coverage was between 2.67 (sample H26 exo -) and 5.33 (sample H27 exo B). In a second sequencing run, the coverage was more homogeneous between DNA samples with variations of 3.26-3.97 between minimal and maximal coverage, a minimum coverage of 936 and a maximum of 11,707 (SI Figure 3). In conclusion, the read coverage of the mitochondrial genome was complete and unbiased in our tested conditions.

**Figure 3:**
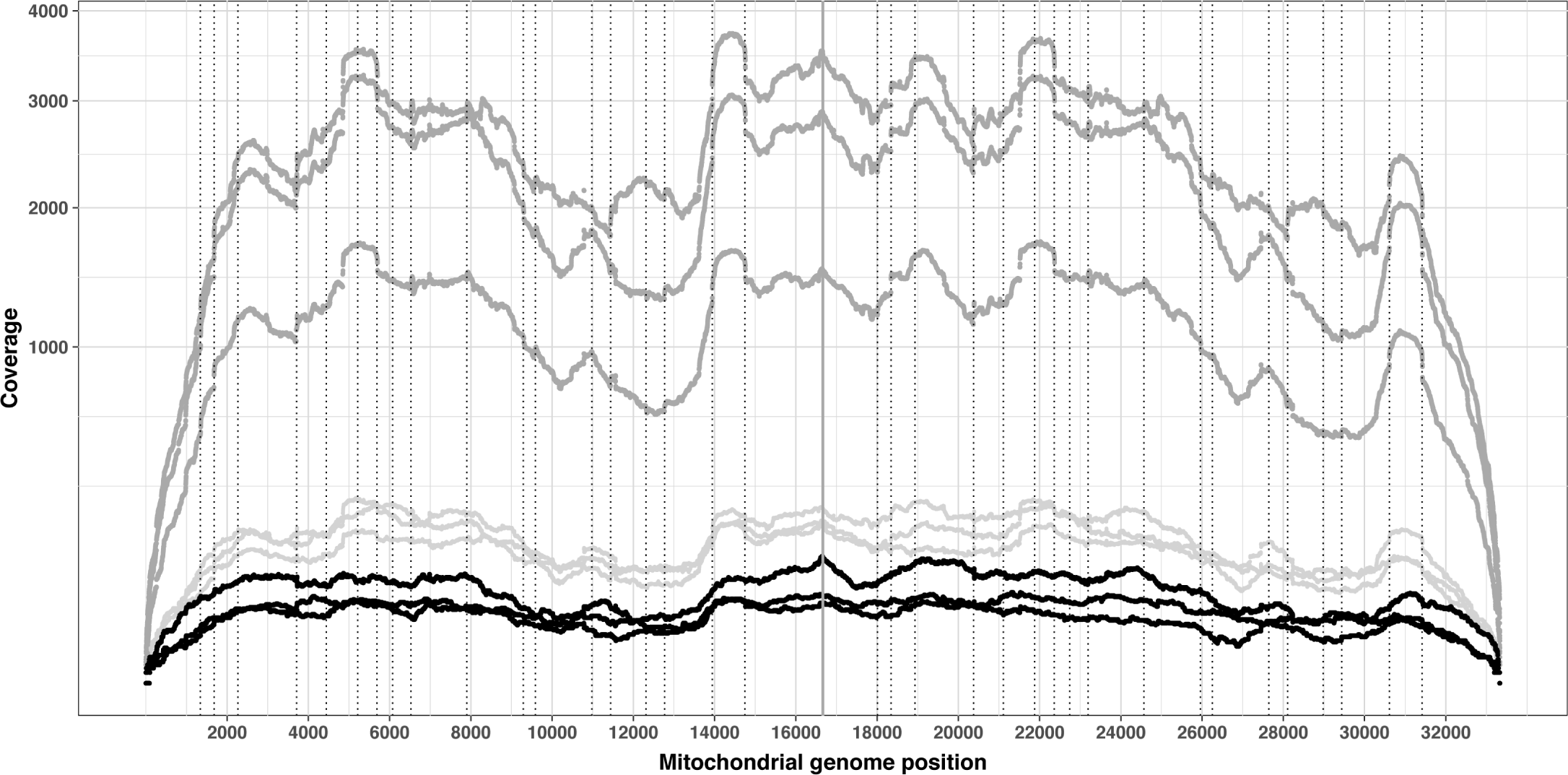
Sequence coverage of the mitochondrial genome. The base position coverage is shown on the y axis. Sequences derived from the exo minus samples are shown in light grey, exo plus A are in dark grey and exo plus B are in black. Coverage was computed from mapping sequences to a duplicated mitochondrial genome. The grey vertical line at position 16666 represents the end of the reference mtDNA sequence. Dotted vertical lines indicate the starting positions of the primers used for MDA.

### Long-read distribution and coverage of the mitochondrial genome

Since we obtained a good coverage of the mitochondrial genome, we wondered about the proportion of long-reads covering a significant portion of the mitochondrial DNA. In H25, H26 and H27 exo A samples, we found respectively 52, 144 and 116 reads longer than half the mitochondrial genome while we only found between one and seven such reads in the other samples (Figure 4, SI table 1). By themselves, these long-reads covered the three horse’s mitochondrial genome at 30.8X, 87.8X and 70.1X (Figure 4B). In about 10 % of these long-reads, we observed that there were multiple alignments between the read and the mitochondrial genome because the longest alignment stretch was smaller than the read length even though we selected reads for which the alignment coverage on mtDNA exceeded 80% (Figure 4A). These cases stemmed from the circular nature of mtDNA and from peculiar reads. For example, we obtained five reads longer than the mitochondrial genome, one of which measuring 17,501 nt and spanning more than one full turn with about 1 kbp of additional coverage (Figure 4B). In other instances, reads were composed of two inverted repeated sequences, possibly originating from the sequencing of both strands of the same molecule, one after the other like in the 2D or 1D2 approaches developed by ONT. In conclusion, we obtained very long-reads from the exonuclease treated DNA spanning large portions of the mitochondrial genome up to its entirety.

**Figure 4:**
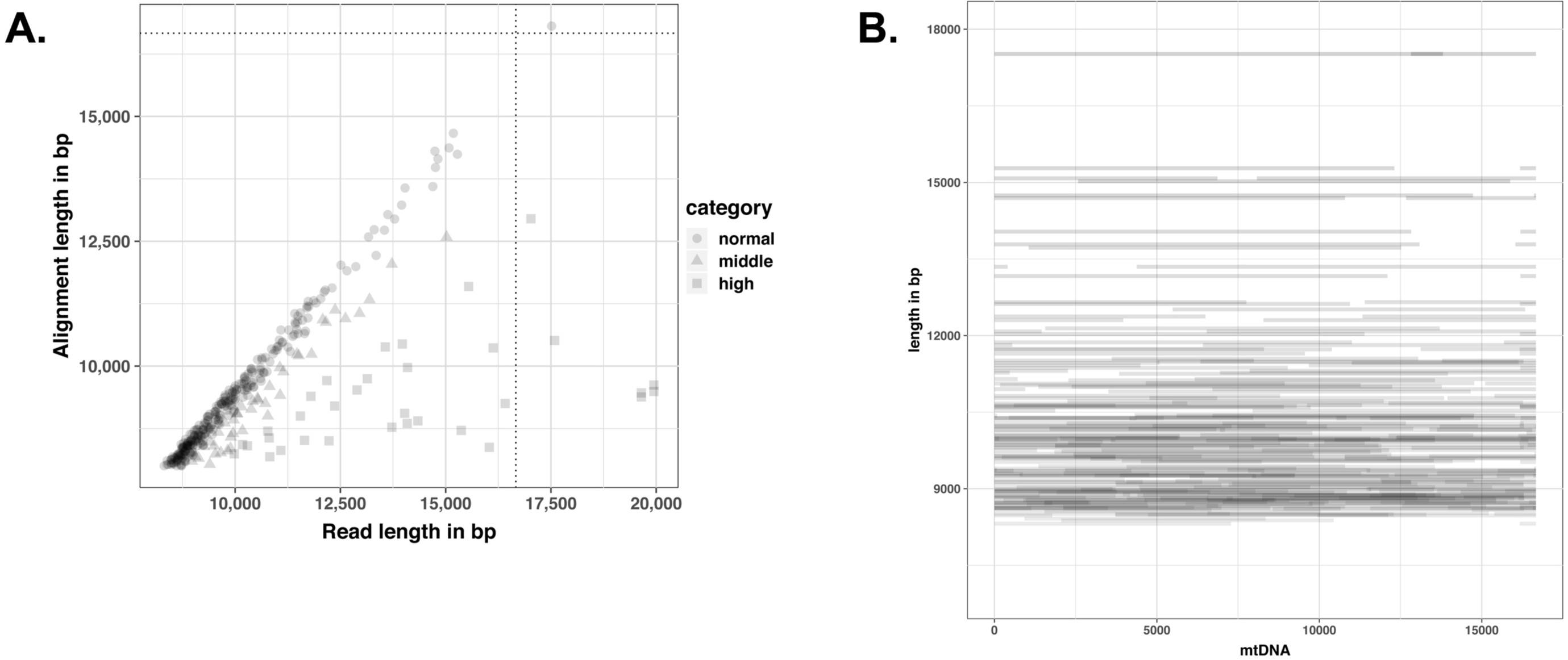
Mitochondrial long-reads. A.This scatter plot shows read and alignment length for reads aligning to more than 8000 bp of mtDNA reference sequence and for which more than 80 % of the read aligns to mtDNA. Dotted lines represent the size of the mitochondrial genome. The different categories correspond to the ratio of read length over alignment length : normal for values less than 1.1, middle for values between 1.1 and 1.2, high for values over 1.2. Values higher than 1 mean that there are mutiple alignments between the read and the mitochondrial genome. B. Long-read coverage display on the horse mitochondrial genome.

These results of the first sequencing run prompted us to reproduce in a second set of experiments the sequencing of the same horse DNAs prepared according to the exo A conditions. We changed the primer pool composition, excluding three primers matching to numerous NUMTs loci in the nuclear genome, performed the exonuclease treatment in duplicate and made a technical replicate of each experiment. We multiplexed the 12 resulting samples for sequencing on the same flow cell. We obtained more reads from this second sequencing run and therefore a much higher coverage of the mitochondrial genome (SI table 1). The fraction of mitochondrial reads ranged from 16 to 20%, with the exception of two technical replicates reaching 50 %. Like previously, the mitochondrial coverage was unbiased (SI Figure 3). The mitochondrial genome coverage by long-reads (> 8000 nt) varied between 118X and 488X, and because the sequencing yield by sample was higher, we also obtained significant coverage by reads longer than 16 kb, from 2 to 40X (SI Table 1, SI Figure 4). In conclusion, we obtained reproducible results in terms of mitochondrial reads enrichment, mitochondrial genome coverage and long-reads.

### Mitochondrial DNA variants calling

We then performed variants calling on all sequencing data sets obtained from three different horses DNA, i.e from the seven sequencing libraries (three from the first sequencing experiment and four from the second sequencing experiment). For this purpose, we used sequence reads of 2000 nt or longer and satisfying a minimal quality threshold of 10, since we obtained a deep coverage and we wanted to avoid reads potentially derived from NUMTs. We used the medaka variant calling method and identified a total of 79 variable positions for H25 samples, 10 for H26 and 84 for H27 (Figure 5). All variants were transitions, and all were found in at least one known horse mtDNA sequence available in GenBank. The majority of these variable positions were identified in the seven sequencing libraries prepared from the same DNA: 70/79 for H25 (88.6%), 10/10 for H26 (100.0%) and 71/84 for H27 (84.5%). The remaining positions were usually of low quality and were counted as false negatives, since they were not identified in at least one sequence sample (see below).

**Figure 5:**
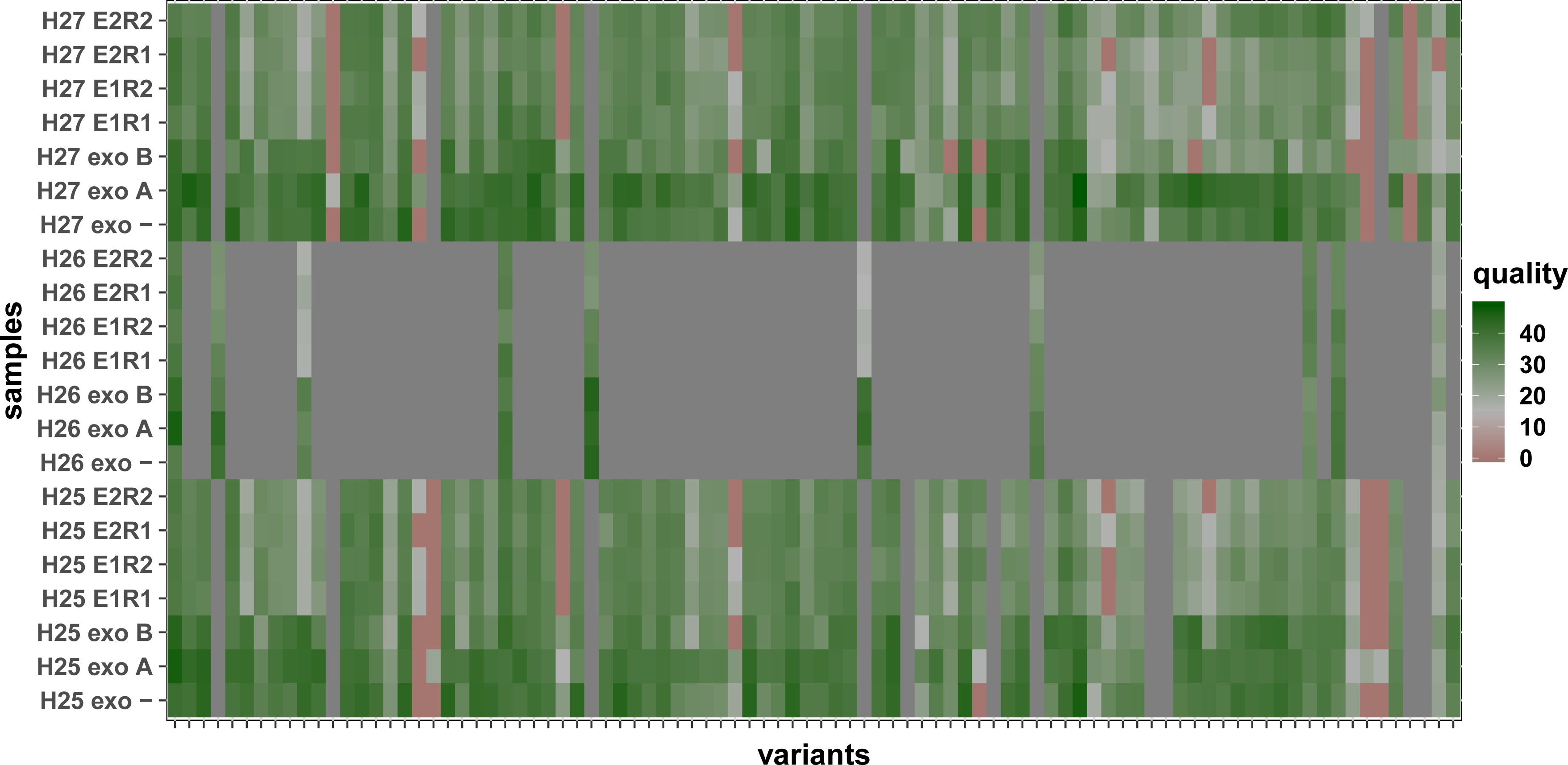
Variant calling from long mtDNA reads. This heatmap shows the different variant identified along the x axis, for the different samples on the y axis. Variants are coloured according to their quality score or are shown in grey when they are not found in a given horse DNA.

We evaluated the validity of these polymorphisms by sequencing nine PCR fragments using Sanger sequencing as ground truth. We could thus compare 7493 bp read using Sanger and Nanopore sequencing. The overall sequence identity between the Sanger and the Nanopore sequences was 99.89%. The nine PCR fragments encompassed 60 variants identified using Nanopore sequencing that enabled a concordance analysis. The F1-score was 0.918 for H25, 1.000 for H26 and 0.911 for H27. Overall we found 52 true positive variants (87%), nine false negative (15%) and one false positive (2%) so that the overall variant recall rate was 0.852, the precision rate was 0.981 and the F1-score was 0.912. We looked at each sample individually to further analyse the between sample variation of true positive and false negative variant calling (SI Table 2). Overall, the mean true positive rate was 94.80% and the false negative rate was 5.20%. The case of H26 was notable since the seven polymorphisms checked were always true positives. The only variation that was always missed by Nanopore sequences was a short deletion in the control region at position 16402 in an array of cytosines. In conclusion, the concordance for SNP calling was very good and depended mostly on variations in the mtDNA control region, a notorious difficult region to sequence.

## Conclusions

We combined two previously described approaches to enrich the fraction of sequence reads derived from mtDNA, namely nuclear DNA removal using an exonuclease treatment and amplification using MDA (33,34). We found that these two techniques require optimisation steps to obtain a significant mtDNA selective amplification factor. For nuclear DNA removal, the exonuclease V amount and incubation time are likely to be dependent on the source and the quality of the input DNA. Using this step, we obtained a 3,000 fold enrichment of the mtDNA/nuDNA ratio, so that theoretically the relative mass amount of mtDNA in the whole cellular DNA increased from 0.63 % (considering 1000 mitochondrial genome per cell) to 99.99 %. This value is in line with the report of Jayaprakash and colleagues (33).

We obtained on the order of a 100,000 fold enrichment after MDA on DNA untreated by exonuclease V, i.e. about ten times more than the value of 100 to 10,000 reported by Marquis and colleagues (34). We used the same reagents as Marquis and colleagues, with the exceptions of primers and DNA, and the two primer concentrations we used in REPLI-g lead to similar enrichment. This difference of a factor of ten may be explained by a difference between human and horse mtDNA/nuDNA ratio, by the qPCR quantification method and by the DNA isolation procedure, the latter being known to impact the quantification of mtDNA copy number (35).

A concern related to MDA is its propensity to produce chimeric molecules, an artefact found in 3 to 6 % of reads derived from Illumina sequencing (36,37). We think that it is unlikely that chimeric reads affect variant calling since chimerism frequency is more than ten times lower than the prevalence of variants. Moreover, such chimeric reads were found to be mostly formed during Illumina sequencing library constructions (36).

We obtained a ratio of mtDNA to nuDNA reads of up to 59%, but this varied for the same starting DNA between different MDA. Still, we obtained a minimum of 16% of reads derived from mtDNA, and this provided an even coverage of the mitochondrial genome. These results are similar to what is reported using some commercial solutions but less than those obtained using PCR on mtDNA extracted using a plasmid preparation kit (38). The principal difference here lies in the horse NUMTs, because most reads mapped to nuDNA originated from NUMTs.

We used a set of primer to amplify horse mtDNA that were uniformly distributed along the mitochondrial genome, and we did not observe a significant effect on the coverage. The absence of a strong coverage bias lead by mtDNA amplification using MDA was also found by Marquis and collaborators (34). In the future, it could be desirable to reduce the number of primers because this could enable the production of larger DNA molecules obtained from MDA. Alternatively, random primers could be used for de novo mtDNA sequencing like in the work of Simison and colleagues (39).

We obtained a sequencing depth of at least 1000X that should allow a much higher level of multiplexing. At the time of writing, multiplexing by ligation is limited to 24 but this is likely to increase in the future since it is only a matter of barcode optimization. According to Quail and colleagues, heterozygous variant calling with a 15x coverage using Ion Torrent or various Illumina datasets was comparable to a coverage of 190x for PacBio datasets (40). We can deduce conservatively that a similar coverage using Nanopore datasets would be enough to call mitochondrial variants in an efficient manner. This means that we could multiplex four or five times more samples, i.e. a hundred, in a single run. With such a capacity, this method could become very competitive in terms of rapidity, scalability and cost-effectiveness.

We estimated at 99.89 % the precision of the obtained sequence, which is satisfactory given that the read median identity was estimated at ~89% for 1D sequencing using R9.0 chemistry. The process of MDA is known to be highly accurate and can not be considered as the limiting factor when it comes to long-read sequencing. But at the same time we observed a variant false negative rate of 5.20% that stems from nanopore sequencing imprecisions around homopolymers of small sizes. The precision is likely to increase through improvement of the sequencing chemistry, like the recent introduction of the R10 flow-cells that combines different pores. We detected only SNPs using our workflow, and the current level of sequencing precision on R9.4 flowcells is likely to preclude the identification of small indels. Further development of variant calling pipelines may alleviate this limitation, and specific tools to detect structural variants could leverage on the added value of long-reads. Another limitation is that the medaka pipeline is tailored for diploid genomes, but suitable models for variant calling on haploid such as FreeBayes could be used.

Previous studies have reported the use of Nanopore sequencing to sequence mitochondrial genomes, usually by sequencing long amplicons (41,42). In a recent study, Lindberg and colleagues evaluated variant allele frequencies from single or mixed DNA sources using both illumina and nanopore sequencing of two overlapping amplicons. In comparison to our findings, they reported a higher precision and recall for single DNA samples (43). Similarly, the added value of long-read sequencing of amplicons for sorting out complex heteroplasmic conditions was leveraged by White and colleagues (41). Only few reports mention in some details the prevalence of mtDNA reads in whole genome shotgun nanopore sequencing. In their report on Nanopore DNA sequencing aboard the International Space Station, Castro-Wallace and colleagues found that 29 out of 83,332 reads (0.03%) from mouse DNA were derived from the mouse mtDNA (44). In another study, Zascavage and colleagues relied on Nanopore long-read sequencing for mtDNA analysis (45). They reported an average coverage of mtDNA reads that varied between 15 and 118 fold from a whole genome sequencing analysis. In their study, the lower coverage limit was inadequate for accurate variant calling, and the coverage range was too large to warrant a routine use.

The identification of causal mtDNA polymorphisms require accurate variant discovery methods. The capacity to phase mtDNA variants, to identify structural variations and nucleotide modifications is a major asset of Nanopore long-read sequencing. The possibility to do so rapidly and at a relatively modest cost is of added interest, and may be of interest for field studies since the MinION sequencer is portable and able to perform real-time sequencing and base-calling (46). At present, the technology is still hindered by the limited level of sequence accuracy but this is rapidly changing as sequencing chemistry and base-calling algorithms are evolving (47,48). The variety of flow cells and equipment available enables to better tailor the sequencing tools according to the number of samples. We believe that this methodology is easily amenable to sequence circular DNA molecules of similar sizes and even beyond, such as plasts, viral circular DNA or endogenous circular DNA (49).

## Methods

### Nuclear DNA depletion

We used DNA samples obtained in the course of a previous project and kindly provided by Dr Céline Robert (50). The DNA were initially extracted from blood samples of H25 (female, Selle Français), H26 (female, Thoroughbred) and H27 (male, Selle Français). We tested nuclear DNA removal by starting with 1 µg of total DNA using either 10 or 20 Units of Exonuclease V (New England Biolabs; UK) and an incubation of 30 minutes or 2 hours at 37°C. We performed the enzymatic digestions in 30 µL reactions containing 1 mM ATP in NEB Buffer 4. We stopped the reaction by adding 0.8 µl of 0.5 M EDTA followed by heat inactivation at 70°C for 30 minutes. DNA was recovered by precipitation with 4 µl of 3M NaCl, 2 µl of 5 mg/ml glycogen and 150 µl of 100 % ethanol. DNA was finally resuspended in 40 µl of nuclease-free water and stored at −20°C until used.

### Whole mitochondrial genome amplification (WMGA)

We used the REPLI-g mitochondrial DNA kit (Qiagen) to perform WMGA according to the manufacturer’s instruction with the following modifications. We designed 18 primers using CLC Genomic Workbench v6 (CLC bio) based on a horse mitochondrial genome (GenBank KX669268.1; see table 1). We adopted the following recommendations for primer design: a length of 10-14 bases, the same number of primers hybridize to each strand; primer hybridization sites are evenly spaced and primers include phosphorothioate modifications between the last three bases of their 3’ end. All primers were purchased from Eurofins Genomics. We tested two primer concentrations: 1 µM and 0.2 µM of each oligonucleotide. The reaction set-up was as follows: 4 µl of DNA treated with Exonuclease V was loaded into a 0.2 ml PCR tube containing 16 µl of RNase DNase free water. 27 µl of REPLI-g mt reaction buffer and 10 µl of horse mitochondrial primer mix (1 µM or 0.2 µM each primer) was added to the DNA and the mixture was incubated at 75°C for 5 min. We added 1 µl of REPLI-g Midi polymerase to the DNA and incubated at 33°C for 8 hours. Inactivation was done at 65 °C for 3 min. The concentration of amplified DNA was quantitated with Quant-iT™ PicoGreen dsDNA reagent (Molecular Probes) on an Infinite M200 Pro (Tecan). We followed published recommendations to avoid possible contaminations (51).

### qPCR experiments

We used qPCR to estimate the ratio of mtDNA over nuclear DNA (nuDNA). The amount of mtDNA was evaluated using three amplicons from the Cox1, Cox3 and Nd2 genes. We also used three nuclear gene loci (Epo, Myod1 and Gapdh) to quantify nuDNA amounts. All primers are given in SI Table 3. As recommended in the REPLI-g mitochondrial DNA kit handbook, the amplified DNA was diluted 1:1000 prior to qPCR. We made two technical replicates for each biological sample, and the same qPCR run contained all samples for a given gene to minimize inter-experimental variations. The 20 µL qPCR reactions were assembled manually. Reactions included 5 µL diluted DNA, 10 µL 2X powered Sybrgreen PCR master mix (Applied Biosystems) and 5 µL primers mix (300 nM each primer). Reactions were incubated in a 96 well optical plate (Applied Biosystems) at 95°C for 10 min, followed by 40 cycles at 95°C for 15 s and 60°C for 1 min in a QuantStudio 12K Flex instrument (Applied Biosystems).

The qPCR data were analysed using the QuantStudio 12K Flex software (Applied Biosystems). Cq values were obtained using the auto baseline option and the threshold was manually set to the same value of ΔRn=0.1 for all amplicons. The Cq values were considered valid if the duplicate measures differed by less than 0.5 cycles, undetermined values were set equivalent to a Ct value of 40. We computed the geometric mean of the Cq values of the three mtDNA and nuDNA amplicons, and then we computed the corresponding fold changes. Formation of primer-dimer and amplification specificity were assessed by melting curve analysis.

### Long-read Sequencing library preparation

Prior to the library construction, we purified the input DNA samples using a 0.5 volume ratio of Agencourt^®^ AMPure^®^ XP magnetic beads. We quantified DNA using a Qubit dsDNA BR assay and performed a debranching treatment on 8-10 µg of DNA in 100 µL reactions containing 50 Units of endonuclease T7E1 (New England Biolabs) in 1X NEB buffer 2, with an incubation of two hours at 37°C. We purified the debranched DNA using one volume of Agencourt^®^ AMPure^®^ XP magnetic beads and checked DNA concentration using a Qubit dsDNA BR assay and DNA integrity of using electrophoresis on a 12K DNA chip (Experion, Bio-Rad).

These DNA samples were then prepared for sequencing according to native barcoding expansion (kit EXP-NBD-103, ONT) and the 1D Native barcoding genomic DNA protocol (kit SQK-LSK108 for the S1 run and SQK-LSK109 for the S2 run, ONT) with some modifications. We treated 2 µg debranched DNA using the UltraII end repair/dA-tailing enzyme mix (New England Biolabs) followed by a purification using a 0.6 volume ratio of AMPure beads. Native barcodes were added by ligation using the Blunt/TA ligase master mix (New England Biolabs) and the ligation reaction was purified using 1 volume ratio of AMPure beads. Equimolar amounts of each barcoded sample were pooled before adapter ligation using the NEBNext Quick ligation Module (New England Biolabs) and a final purification using a 0.4 volume ratio of AMPure beads with the S Fragment buffer.

For the S1 sequencing run, we used DNA amplified with the 18 primers and multiplexed nine samples using barcodes one to nine. We loaded 195 ng of sequencing library. In this S1 run, each of the three horse’s DNA libraries was derived from three different conditions of exonuclease and amplification, so that each horse’s DNA was independently sequenced three times. For the S2 sequencing run, we used DNA amplified with 15 primers, excluding three that were found to match NUMTs loci, and multiplexed twelve samples using barcodes one to twelve. We loaded 439 ng of the sequencing library. In this S2 run, we performed each exonuclease treatment in duplicate and each amplification in duplicate, so that each horse’s DNA was independently sequenced four times. Altogether, each horse’s DNA was sequenced across seven libraries.

We performed the sequencing using R9.4 flowcells from two different batches on a MinION sequencer (ONT) operated using MinION Release v18.05.5 for the S1 run and 19.05.0 for the S2 run. The MinION sequencing device was connected to a computer running the linux operating system Ubuntu 16.04 LTS.

### Data analysis

We used guppy for basecalling with the flip-flop model, trimming and homopolymer correction (guppy-cpu-v2.3.1). We then used guppy_barcoder for demultiplexing. For variant calling, we selected reads longer than 2000 bp with a mean quality higher than Q10 using NanoFilt (52). The filtered sequences were then aligned using minimap2 on the *Equus caballus* genome reference assembly (GCF_002863925.1_EquCab3.0) (53). We substituted the mitochondrial genome present in this assembly by a duplicated, pseudo-circularized, mitochondrial genome (GenBank JN398434) to increase the recovery of mapping to the circular mitochondrial genome. We used bioawk and samtools to extract mitochondrial reads and compute their quality scores (54). These mtDNA reads were aligned with minimap2 on a reference horse mtDNA (JN398434) to call variants using the medaka pipeline, and filtered using a quality cut-off of 15. All visualisation scripts were written using the ggplot2 package in R3.6.0 (55,56).

### Sanger Sequencing for SNP validation

We designed nine primer pairs to amplify nine fragments evenly spaced along the horse mtDNA (see table 1). PCRs were performed on 4 ng of the same DNA used for long-read sequencing and using the 2X HotStartTaq Master Mix (Qiagen). PCR products were checked by agarose gel electrophoresis and purified using Nucleospin Gel and PCR clean-up kit (Macherey-Nagel). Cycle sequencing reactions were performed using the BigDye 3.1 chemistry, purified by precipitation and sequenced using an ABI 3130 sequencer (Applied Biosystems). Base calling was performed using the Sequencing analysis software version 5. 3 (Applied Biosystems).

## Declarations

### Ethics approval and consent to participate

Horse’s DNA samples were obtained in the course of a previous project (50).

## Consent for publication

Not applicable

## Availability of data and materials

The datasets generated and/or analysed during the current study are available at the European Nucleotide Archive PRJEB35937

## Competing Interests

The authors declare no competing interests.

## Funding

### Author’s contributions

S.D-P, E.B., N.P. designed the experiment.

S.D-P. performed the DNA pre-treatments (Exonuclease treatment and MDA), qPCR analyses, PCR amplifications in S.D. and E.B. laboratory.

S.D-P. and N.P. performed sequencing library preparations in N.P. laboratory.

N.P. performed the Sanger sequencing experiments.

N.P. carried out the bioinformatics analyses.

N.P. drafted the main text and prepared figures with inputs from S.D-P and E.B. All authors reviewed the manuscript and approved the final version.

## Acknowledgements

The authors would like to thank Prof. Céline Robert for the kind gift of horse’s DNA.

## Supplementary information

**SI_table 1 : Sequencing statistics, coverage, long-reads**

**SI_table 2 : Sanger resequencing informations**

**SI_table 3 : Primers**

**SI_Figure 1 : DNA quantity and fragment size after REPLI-g**

**SI_Figure 2 : Distribution of raw read length and quality by barcode**

**SI_Figure 3 : Proportions of mtDNA and nuDNA sequences and sequence coverage in a second sequencing run**

**SI_Figure 4 : Distribution of long reads in a second sequencing run**

## References

1. Roger AJ, Muñoz-Gómez SA, Kamikawa R. The Origin and Diversification of Mitochondria. Curr Biol CB. 6 nov 2017;27(21):R1177–92.

2. Gray MW. Mitochondrial evolution. Cold Spring Harb Perspect Biol. 1 sept 2012;4(9):a011403.

3. Nass MM, Nass S. Intramitochondrial fibers with DNA characteristics. I. Fixation and electron staining reactions. J Cell Biol. déc 1963;19:593–611.

4. Smith DR. The past, present and future of mitochondrial genomics: have we sequenced enough mtDNAs? Brief Funct Genomics. janv 2016;15(1):47–54.

5. Kogelnik AM, Lott MT, Brown MD, Navathe SB, Wallace DC. MITOMAP: a human mitochondrial genome database. Nucleic Acids Res. 1 janv 1996;24(1):177–9.

6. Bernt M, Braband A, Schierwater B, Stadler PF. Genetic aspects of mitochondrial genome evolution. Mol Phylogenet Evol. 1 nov 2013;69(2):328–38.

7. Ballard JWO, Whitlock MC. The incomplete natural history of mitochondria. Mol Ecol. avr 2004;13(4):729–44.

8. Allio R, Donega S, Galtier N, Nabholz B. Large Variation in the Ratio of Mitochondrial to Nuclear Mutation Rate across Animals: Implications for Genetic Diversity and the Use of Mitochondrial DNA as a Molecular Marker. Mol Biol Evol. 1 nov 2017;34(11):2762–72.

9. Ramos A, Santos C, Mateiu L, Gonzalez M del M, Alvarez L, Azevedo L, et al. Frequency and pattern of heteroplasmy in the complete human mitochondrial genome. PloS One. 2013;8(10):e74636.

10. Pamilo P, Viljakainen L, Vihavainen A. Exceptionally high density of NUMTs in the honeybee genome. Mol Biol Evol. juin 2007;24(6):1340–6.

11. Ricchetti M, Tekaia F, Dujon B. Continued colonization of the human genome by mitochondrial DNA. PLoS Biol. sept 2004;2(9):E273.

12. Nergadze SG, Lupotto M, Pellanda P, Santagostino M, Vitelli V, Giulotto E. Mitochondrial DNA insertions in the nuclear horse genome. Anim Genet. déc 2010;41 Suppl 2:176–85.

13. Dayama G, Emery SB, Kidd JM, Mills RE. The genomic landscape of polymorphic human nuclear mitochondrial insertions. Nucleic Acids Res. 10 nov 2014;42(20):12640–9.

14. Robin ED, Wong R. Mitochondrial DNA molecules and virtual number of mitochondria per cell in mammalian cells. J Cell Physiol. sept 1988;136(3):507–13.

15. Bensasson D, Zhang D-X, Hartl DL, Hewitt GM. Mitochondrial pseudogenes: evolution’s misplaced witnesses. Trends Ecol Evol. 1 juin 2001;16(6):314–21.

16. Nguyen TTT, Murphy NP, Austin CM. Amplification of multiple copies of mitochondrial Cytochrome b gene fragments in the Australian freshwater crayfish, Cherax destructor Clark (Parastacidae: Decapoda). Anim Genet. août 2002;33(4):304–8.

17. Yao Y-G, Kong Q-P, Salas A, Bandelt H-J. Pseudomitochondrial genome haunts disease studies. J Med Genet. déc 2008;45(12):769–72.

18. Maddock ST, Briscoe AG, Wilkinson M, Waeschenbach A, Mauro DS, Day JJ, et al. Next-Generation Mitogenomics: A Comparison of Approaches Applied to Caecilian Amphibian Phylogeny. PLOS ONE. 9 juin 2016;11(6):e0156757.

19. Briscoe AG, Hopkins KP, Waeschenbach A. High-Throughput Sequencing of Complete Mitochondrial Genomes. Methods Mol Biol Clifton NJ. 2016;1452:45–64.

20. Istace B, Friedrich A, d’Agata L, Faye S, Payen E, Beluche O, et al. de novo assembly and population genomic survey of natural yeast isolates with the Oxford Nanopore MinION sequencer. GigaScience [Internet]. 1 févr 2017 [cité 18 déc 2019];6(2). Disponible sur: http://academic.oup.com/gigascience/article/6/2/giw018/2865217

21. Ranjard L, Wong TKF, Külheim C, Rodrigo AG, Ragg NLC, Patel S, et al. Complete mitochondrial genome of the green-lipped mussel, Perna canaliculus (Mollusca: Mytiloidea), from long nanopore sequencing reads. Mitochondrial DNA Part B. 2 janv 2018;3(1):175–6.

22. Gan HM, Grandjean F, Jenkins TL, Austin CM. Absence of evidence is not evidence of absence: Nanopore sequencing and complete assembly of the European lobster (Homarus gammarus) mitogenome uncovers the missing nad2 and a new major gene cluster duplication. BMC Genomics. 3 mai 2019;20(1):335.

23. Gan HM, Linton SM, Austin CM. Two reads to rule them all: Nanopore long read-guided assembly of the iconic Christmas Island red crab, Gecarcoidea natalis (Pocock, 1888), mitochondrial genome and the challenges of AT-rich mitogenomes. Mar Genomics. juin 2019;45:64–71.

24. Barrey E, Mucher E, Jeansoule N, Larcher T, Guigand L, Herszberg B, et al. Gene expression profiling in equine polysaccharide storage myopathy revealed inflammation, glycogenesis inhibition, hypoxia and mitochondrial dysfunctions. BMC Vet Res. 7 août 2009;5:29.

25. Barrey E, Jayr L, Mucher E, Gospodnetic S, Joly F, Benech P, et al. Transcriptome analysis of muscle in horses suffering from recurrent exertional rhabdomyolysis revealed energetic pathway alterations and disruption in the cytosolic calcium regulation. Anim Genet. juin 2012;43(3):271–81.

26. Mach N, Plancade S, Pacholewska A, Lecardonnel J, Rivière J, Moroldo M, et al. Integrated mRNA and miRNA expression profiling in blood reveals candidate biomarkers associated with endurance exercise in the horse. Sci Rep. 10 mars 2016;6:22932.

27. Ricard A, Robert C, Blouin C, Baste F, Torquet G, Morgenthaler C, et al. Endurance Exercise Ability in the Horse: A Trait with Complex Polygenic Determinism. Front Genet. 2017;8:89.

28. Keyser-Tracqui C, Blandin-Frappin P, Francfort H-P, Ricaut F-X, Lepetz S, Crubézy E, et al. Mitochondrial DNA analysis of horses recovered from a frozen tomb (Berel site, Kazakhstan, 3rd Century BC). Anim Genet. juin 2005;36(3):203–9.

29. Achilli A, Olivieri A, Soares P, Lancioni H, Hooshiar Kashani B, Perego UA, et al. Mitochondrial genomes from modern horses reveal the major haplogroups that underwent domestication. Proc Natl Acad Sci U S A. 14 févr 2012;109(7):2449–54.

30. Fages A, Hanghøj K, Khan N, Gaunitz C, Seguin-Orlando A, Leonardi M, et al. Tracking Five Millennia of Horse Management with Extensive Ancient Genome Time Series. Cell. 30 mai 2019;177(6):1419–1435.e31.

31. Wolff JN, Shearman DCA, Brooks RC, Ballard JWO. Selective enrichment and sequencing of whole mitochondrial genomes in the presence of nuclear encoded mitochondrial pseudogenes (numts). PloS One. 2012;7(5):e37142.

32. Santibanez-Koref M, Griffin H, Turnbull DM, Chinnery PF, Herbert M, Hudson G. Assessing mitochondrial heteroplasmy using next generation sequencing: A note of caution. Mitochondrion. mai 2019;46:302–6.

33. Jayaprakash AD, Benson EK, Gone S, Liang R, Shim J, Lambertini L, et al. Stable heteroplasmy at the single-cell level is facilitated by intercellular exchange of mtDNA. Nucleic Acids Res. 27 févr 2015;43(4):2177–87.

34. Marquis J, Lefebvre G, Kourmpetis YAI, Kassam M, Ronga F, De Marchi U, et al. MitoRS, a method for high throughput, sensitive, and accurate detection of mitochondrial DNA heteroplasmy. BMC Genomics [Internet]. 26 avr 2017 [cité 4 nov 2019];18. Disponible sur: https://www.ncbi.nlm.nih.gov/pmc/articles/PMC5405551/

35. Fazzini F, Schöpf B, Blatzer M, Coassin S, Hicks AA, Kronenberg F, et al. Plasmid-normalized quantification of relative mitochondrial DNA copy number. Sci Rep. 18 2018;8(1):15347.

36. Ellegaard KM, Klasson L, Andersson SGE. Testing the Reproducibility of Multiple Displacement Amplification on Genomes of Clonal Endosymbiont Populations. PLOS ONE. 27 nov 2013;8(11):e82319.

37. Tu J, Guo J, Li J, Gao S, Yao B, Lu Z. Systematic Characteristic Exploration of the Chimeras Generated in Multiple Displacement Amplification through Next Generation Sequencing Data Reanalysis. PLoS ONE [Internet]. 6 oct 2015 [cité 6 nov 2019];10(10). Disponible sur: https://www.ncbi.nlm.nih.gov/pmc/articles/PMC4595205/

38. Quispe-Tintaya W, White RR, Popov VN, Vijg J, Maslov AY. Fast mitochondrial DNA isolation from mammalian cells for next-generation sequencing. BioTechniques. sept 2013;55(3):133–6.

39. Simison WB, Lindberg DR, Boore JL. Rolling circle amplification of metazoan mitochondrial genomes. Mol Phylogenet Evol. mai 2006;39(2):562–7.

40. Quail MA, Smith M, Coupland P, Otto TD, Harris SR, Connor TR, et al. A tale of three next generation sequencing platforms: comparison of Ion Torrent, Pacific Biosciences and Illumina MiSeq sequencers. BMC Genomics. 24 juill 2012;13(1):341.

41. White EJ, Ross T, Lopez E, Nikiforov A, Gault C, Batorsky R, et al. Chasing a moving target: Detection of mitochondrial heteroplasmy for clinical diagnostics. bioRxiv. 19 nov 2017;222109.

42. Tan AS, Baty JW, Dong L-F, Bezawork-Geleta A, Endaya B, Goodwin J, et al. Mitochondrial Genome Acquisition Restores Respiratory Function and Tumorigenic Potential of Cancer Cells without Mitochondrial DNA. Cell Metab. 6 janv 2015;21(1):81–94.

43. Lindberg MR, Schmedes SE, Hewitt FC, Haas JL, Ternus KL, Kadavy DR, et al. A Comparison and Integration of MiSeq and MinION Platforms for Sequencing Single Source and Mixed Mitochondrial Genomes. PloS One. 2016;11(12):e0167600.

44. Castro-Wallace SL, Chiu CY, John KK, Stahl SE, Rubins KH, McIntyre ABR, et al. Nanopore DNA Sequencing and Genome Assembly on the International Space Station. Sci Rep. 21 déc 2017;7(1):1–12.

45. Zascavage RR, Thorson K, Planz JV. Nanopore sequencing: An enrichment-free alternative to mitochondrial DNA sequencing. Electrophoresis. 2019;40(2):272–80.

46. Loose M, Malla S, Stout M. Real-time selective sequencing using nanopore technology. Nat Methods. 2016;13(9):751–4.

47. Noakes MT, Brinkerhoff H, Laszlo AH, Derrington IM, Langford KW, Mount JW, et al. Increasing the accuracy of nanopore DNA sequencing using a time-varying cross membrane voltage. Nat Biotechnol. 2019;37(6):651–6.

48. Rang FJ, Kloosterman WP, de Ridder J. From squiggle to basepair: computational approaches for improving nanopore sequencing read accuracy. Genome Biol. 13 2018;19(1):90.

49. McNaughton AL, Roberts HE, Bonsall D, Cesare M de, Mokaya J, Lumley SF, et al. Illumina and Nanopore methods for whole genome sequencing of hepatitis B virus (HBV). Sci Rep. 8 mai 2019;9(1):1–14.

50. Robert C, Valette J-P, Jacquet S, Lepeule J, Denoix J-M. Study design for the investigation of likely aetiological factors of juvenile osteochondral conditions (JOCC) in foals and yearlings. Vet J Lond Engl 1997. juill 2013;197(1):36–43.

51. Champlot S, Berthelot C, Pruvost M, Bennett EA, Grange T, Geigl E-M. An efficient multistrategy DNA decontamination procedure of PCR reagents for hypersensitive PCR applications. PloS One. 28 sept 2010;5(9).

52. De Coster W, D’Hert S, Schultz DT, Cruts M, Van Broeckhoven C. NanoPack: visualizing and processing long-read sequencing data. Bioinformatics. 1 août 2018;34(15):2666–9.

53. Li H. Minimap2: pairwise alignment for nucleotide sequences. Bioinforma Oxf Engl. 15 2018;34(18):3094–100.

54. Li H, Handsaker B, Wysoker A, Fennell T, Ruan J, Homer N, et al. The Sequence Alignment/Map format and SAMtools. Bioinforma Oxf Engl. 15 août 2009;25(16):2078–9.

55. Wickham H. ggplot2: Elegant Graphics for Data Analysis [Internet]. New York: Springer-Verlag; 2009 [cité 6 nov 2019]. (Use R!). Disponible sur: https://www.springer.com/gp/book/9780387981413

56. R Core Team. R: A language and environment for statistical computing. [Internet]. 2019. Disponible sur: www.R-project.org

