## Supplementary_information for "A new method for long-read sequencing of animal mitochondrial genomes: application to the identification of equine mitochondrial DNA variants"

### Quantity of DNA after Repli-G

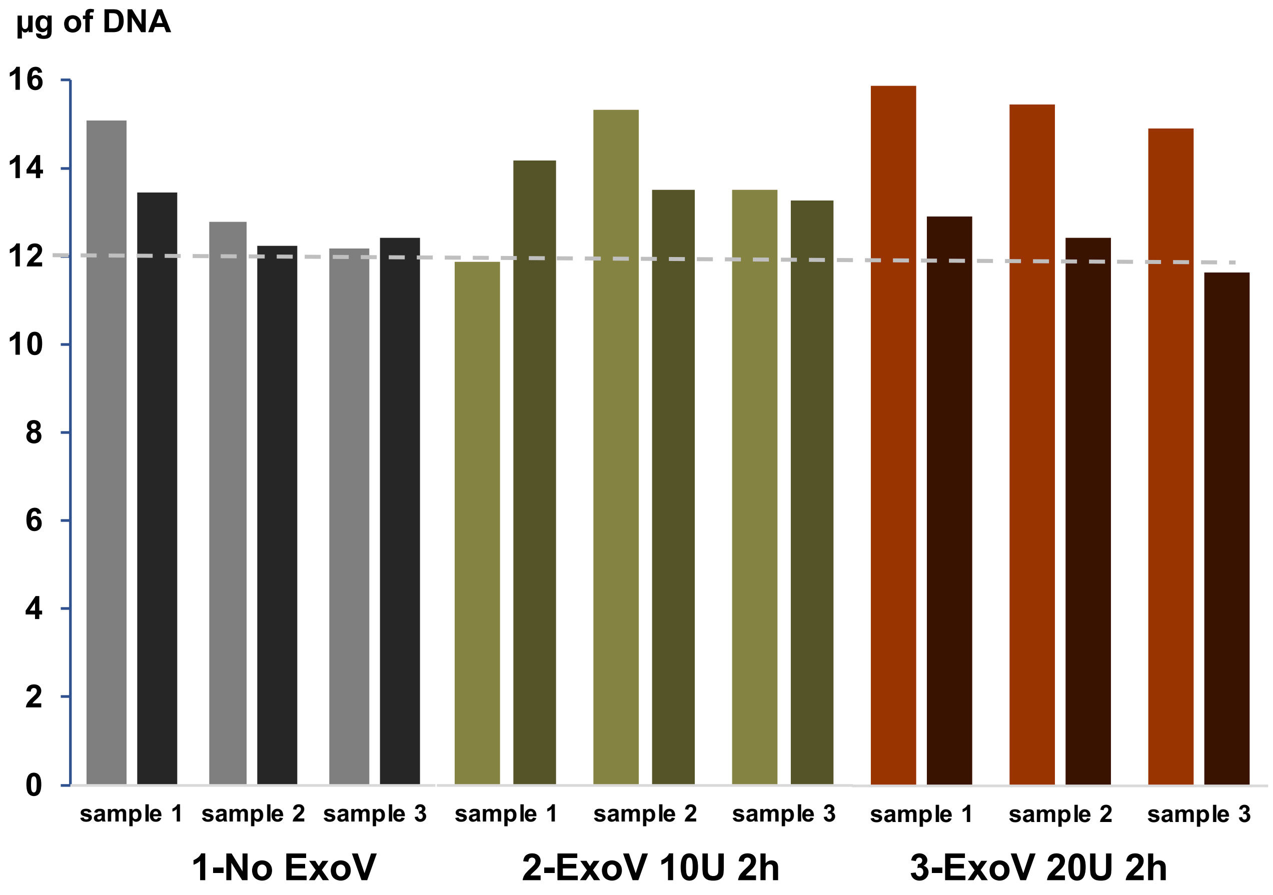


### Fragment size after Repli-G

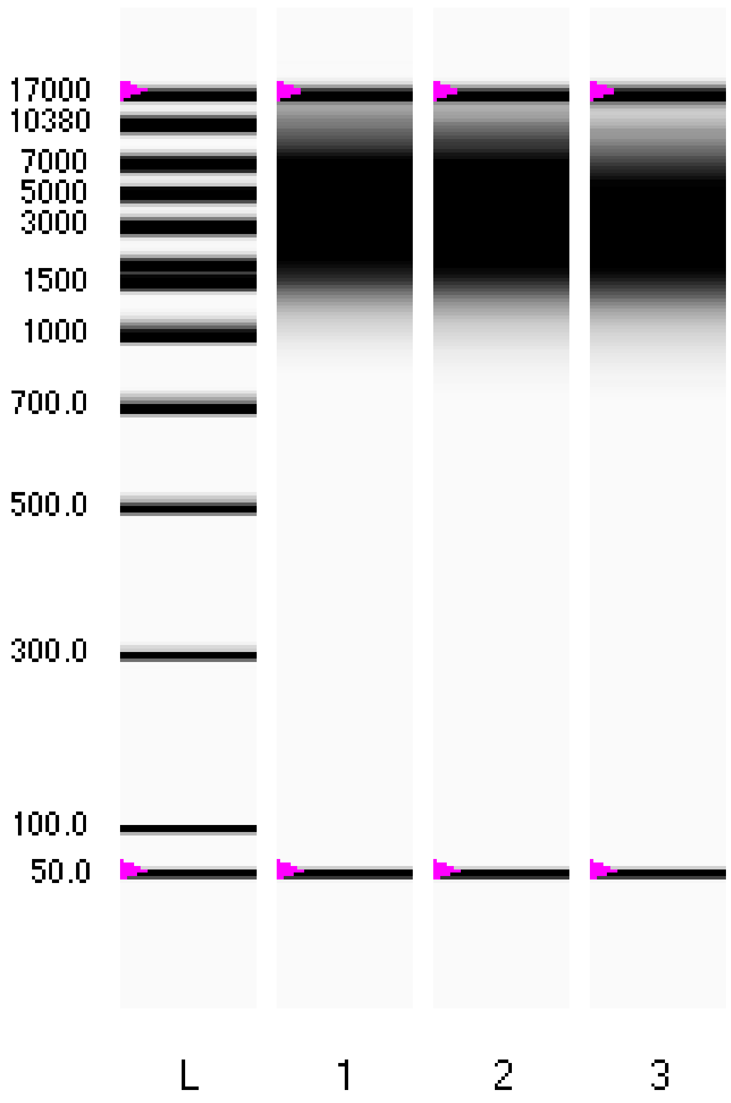


Supplementary data. Dhorne-Pollet et al.

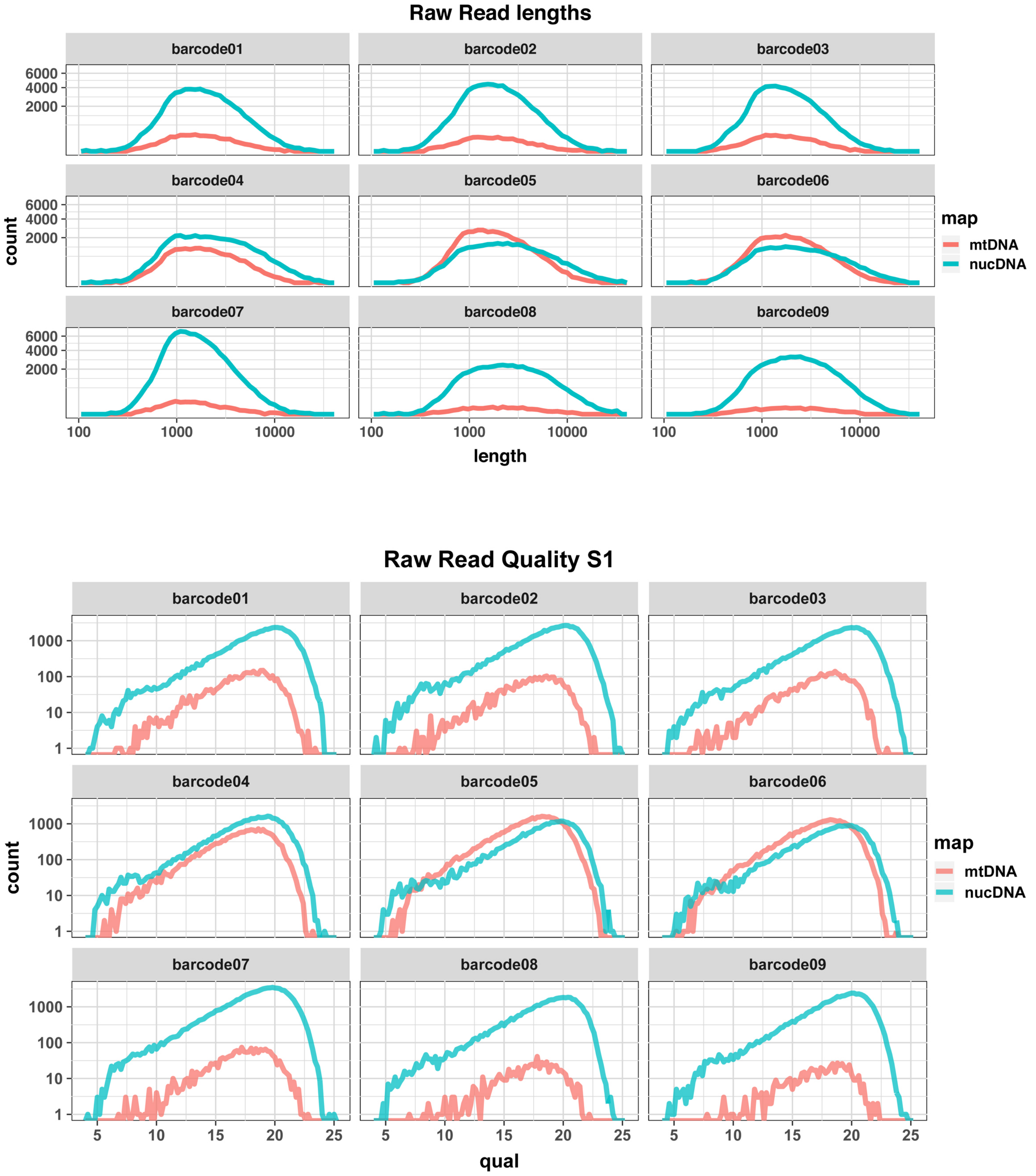
Raw Read lengths
barcode01
barcode02
barcode03
barcode04
barcode05
barcode06
barcode07
barcode08
barcode09
6000
4000
2000
count
map
mtDNA
nucDNA
100
1000
10000
length
Raw Read Quality S1
1000
100
10
1
5
10
15
20
25
qual

SI Figure 2

### a. Proportions of mtDNA and nuDNA sequences

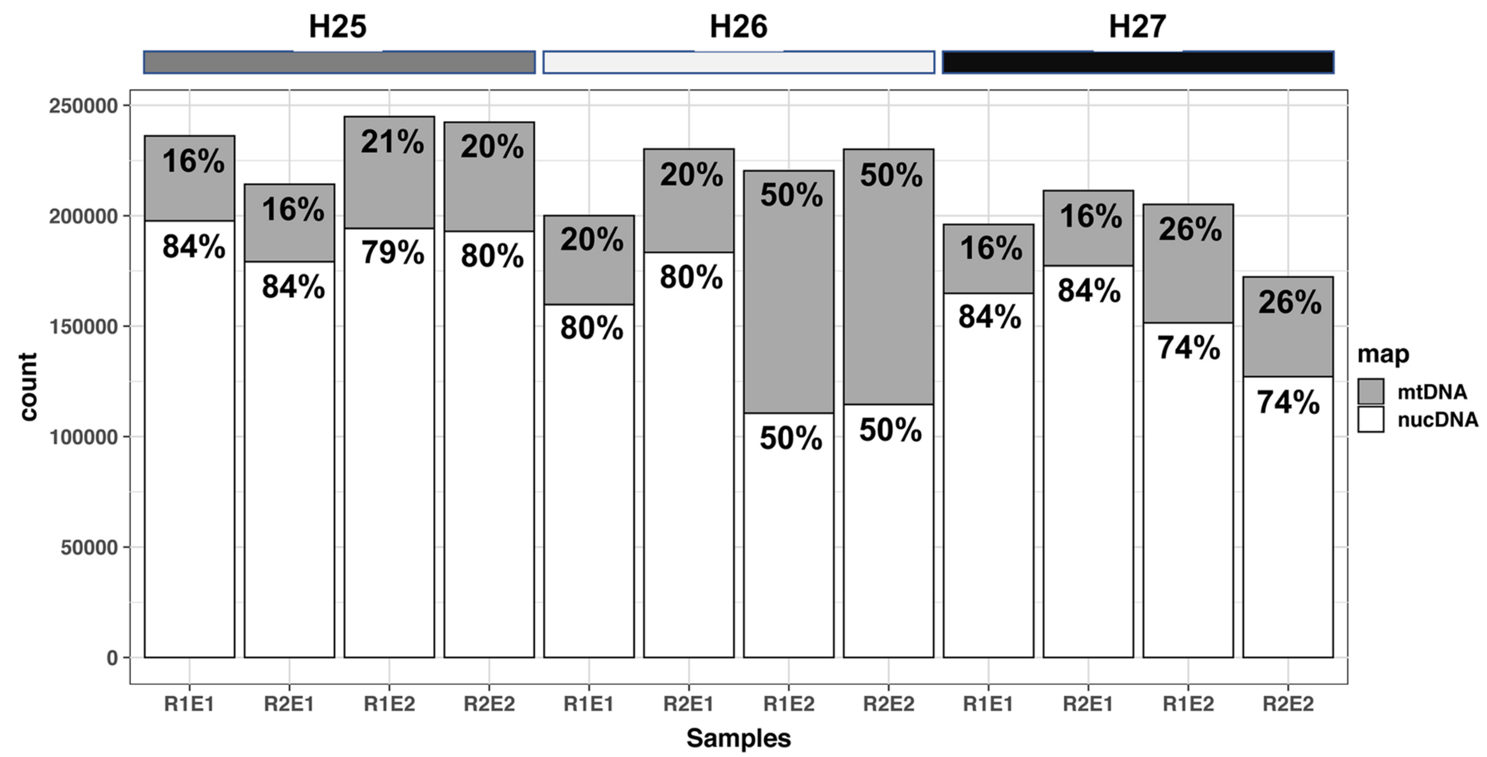


### b. Sequence coverage

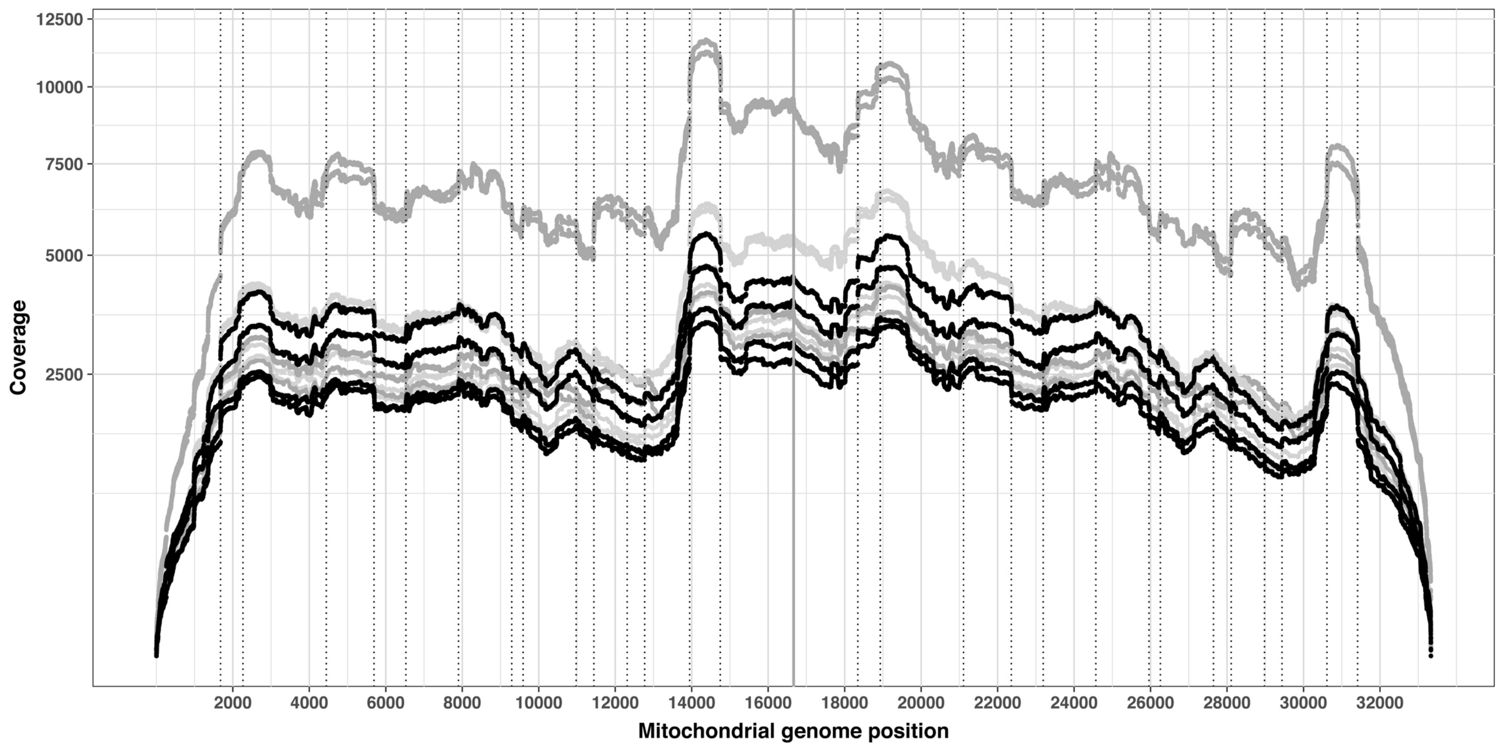


SI Figure 3

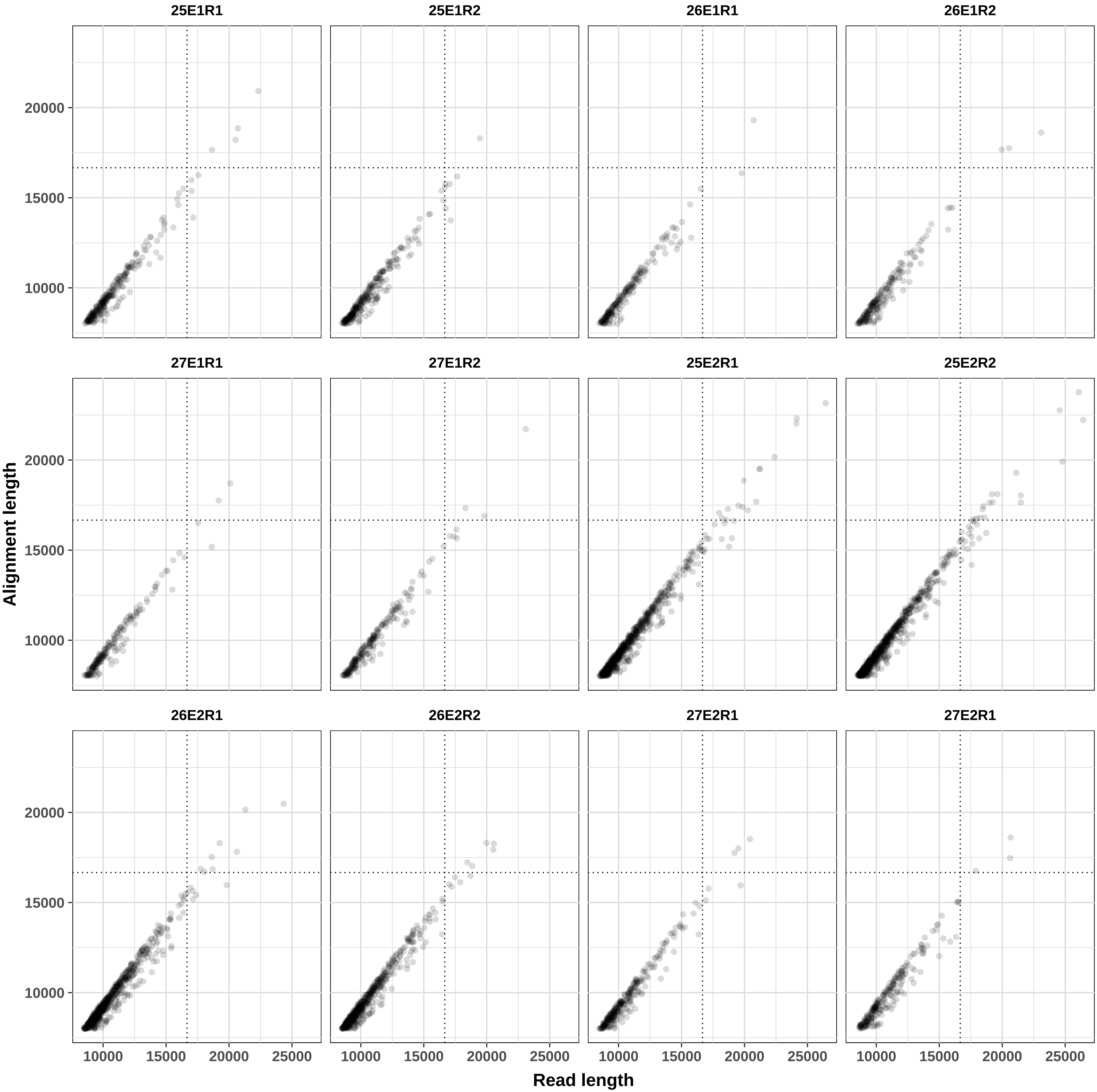

25E1R1
25E1R2
26E1R1
26E1R2
27E1R1
27E1R2
25E2R1
25E2R2
26E2R1
26E2R2
27E2R1
27E2R1
Alignment length
Read length
10000
15000
20000
25000

| Sample<br>Barcode | H25 R1E1<br>Barcode01 | H25 R2E1<br>Barcode02 | H26 R1E1<br>Barcode03 | H26 R2E1<br>Barcode04 | H27 R1E1<br>Barcode05 | H27 R2E1<br>Barcode06 | H25 R1E2<br>Barcode07 | H25 R2E2<br>Barcode08 | H26 R1E2<br>Barcode09 | H26 R2E2<br>Barcode10 | H27 R1E2<br>Barcode11 | H27 R2E2<br>Barcode12 |
|---|---|---|---|---|---|---|---|---|---|---|---|---|
| Nb read length > 8000 nt | 301 | 320 | 228 | 209 | 206 | 252 | 822 | 836 | 725 | 644 | 320 | 276 |
| Nb read length > 8000 nt and coverage > 80% | 261 | 275 | 203 | 191 | 182 | 212 | 717 | 729 | 638 | 552 | 276 | 232 |
| long read mtDNA fold coverage | 170 | 177 | 130 | 122 | 118 | 141 | 479 | 488 | 415 | 353 | 178 | 152 |
| Nb read length > 16666 nt | 16 | 23 | 6 | 7 | 7 | 11 | 57 | 68 | 30 | 27 | 14 | 14 |
| Nb read length > 16666 nt and coverage > 80% | 8 | 7 | 2 | 3 | 4 | 7 | 26 | 35 | 14 | 10 | 6 | 3 |
| very long read mtDNA fold coverage | 9 | 7 | 2 | 4 | 5 | 8 | 31 | 40 | 16 | 11 | 7 | 4 |

| PCR fragment | OL. Start | OL. End | PCR Span | Sanger coverage | Variants in span Sanger | H25-True | H25-False | H26-True | H26-False | H27-True | H27-False | True positive | False negative | False positive | Note | Context | Context |
|---|---|---|---|---|---|---|---|---|---|---|---|---|---|---|---|---|---|
| PCR fragment 1 | 930 | 1929 | 999 | 956 | 5 | 4 | 0 | 1 | 0 | 4 | 0 | 5 | 0 | 0 | | | |
| PCR fragment 2 | 2462 | 3243 | 781 | 763 | 4 | 4 | 0 | 1 | 0 | 4 | 0 | 4 | 0 | 0 | | | |
| PCR fragment 3 | 3440 | 4458 | 1018 | 906 | 4 | 3 | **1** | 0 | 0 | 3 | 1 | 3 | 1 | 0 | **False negative: Sanger C->T at 3941** | AATAAA[C/T]CCCCT | |
| PCR fragment 4 | 7104 | 8085 | 981 | 903 | 5 | 4 | 1 | 1 | 0 | 4 | 1 | 4 | 1 | 0 | **False negative: Sanger T->C at 8057** | TTCCC[T/C]TCACCC | |
| PCR fragment 5 | 7975 | 8937 | 962 | 847 | 6 | 5 | 0 | 1 | 0 | 5 | 0 | 6 | 0 | 0 | | | |
| PCR fragment 6 | 11523 | 12522 | 999 | 972 | 8 | 7 | 0 | 1 | 0 | 7 | 0 | 8 | 0 | 0 | | | |
| PCR fragment 7 | 13315 | 14029 | 714 | 605 | 4 | 2 | 1 | 0 | 0 | 4 | 1 | 4 | 1 | 0 | **False negative: Sanger G->A at 13519** | CACCCA[G/A]AAAGGA | |
| PCR fragment 8 | 14915 | 15789 | 874 | 861 | 13 | 9 | 2 | 0 | 0 | 7 | 3 | 10 | 3 | 0 | **False negative: Sanger G->A at 15492;Sanger GA->AG at 15645** | CAATCC[G->A]CCCT | TACAT[GA/AG]GTACA |
| PCR fragment 9 | 15800 | 16632 | 832 | 680 | 11 | 7 | **3** | 2 | 0 | 8 | **3** | 8 | **3** | 1 | **False negative:Sanger G->A at 16118; TG->CA at 16126; insertion C at 16402** | CATAG[G/A]AAACAAG[TG/CA]CACC | CCCCCC[C]TACCCCCCA |
| All PCR fragments | | | | 7493 | 60 | 45 | 8 | 7 | 0 | 46 | 9 | 52 | 9 | 1 | | | |
| Identity | | | | 99.89% | | | | | | | | 87% | 15% | 2% | | | |

Primers used in this study

| Primers | Application |
|---|---|
| MT-FWD-1 | MDA |
| MT-FWD-3 | MDA |
| MT-FWD-5 | MDA |
| MT-FWD-6 | MDA |
| MT-FWD-7 | MDA |
| MT-FWD-9 | MDA |
| MT-FWD-11 | MDA |
| MT-FWD-12 | MDA |
| MT-FWD-13 | MDA |
| MT-REV-1 | MDA |
| MT-REV-3 | MDA |
| MT-REV-5 | MDA |
| MT-REV-6 | MDA |
| MT-REV-14 | MDA |
| MT-REV-10 | MDA |
| MT-REV-9 | MDA |
| MT-REV-13 | MDA |
| MT-REV-14 | MDA |
| COX1F | qPCR |
| COX1R | qPCR |
| COX3F | qPCR |
| COX3R | qPCR |
| ND2F | qPCR |
| ND2R | qPCR |
| GAPDHF | qPCR |
| GAPDHR | qPCR |
| MYOD1F | qPCR |
| MYOD1R | qPCR |
| EPOF | qPCR |
| EPOR | qPCR |
| mtDNA_primer-1F | PCR/Sequencing |
| mtDNA_primer-1R | PCR/Sequencing |
| mtDNA_primer-2F | PCR/Sequencing |
| mtDNA_primer-2R | PCR/Sequencing |
| mtDNA_primer-3F | PCR/Sequencing |
| mtDNA_primer-3R | PCR/Sequencing |
| mtDNA_primer-4F | PCR/Sequencing |
| mtDNA_primer-4R | PCR/Sequencing |
| mtDNA_primer-5F | PCR/Sequencing |
| mtDNA_primer-5R | PCR/Sequencing |
| mtDNA_primer-6F | PCR/Sequencing |
| mtDNA_primer-6R | PCR/Sequencing |
| mtDNA_primer-7F | PCR/Sequencing |

| | |
|---|---|
| mtDNA_primer-7R | PCR/Sequencing |
| mtDNA_primer-8F | PCR/Sequencing |
| mtDNA_primer-8R | PCR/Sequencing |
| mtDNA_primer-9F | PCR/Sequencing |
| mtDNA_primer-9R | PCR/Sequencing |

| séquences (5' -3') * | Reference sequence |
|---|---|
| CCCAACAATCAACC | KX669268.1 |
| CCCACCACAAACA | KX669268.1 |
| AAAAGGCGGGAGA | KX669268.1 |
| ATGGGGGGATTTGT | KX669268.1 |
| CGACACACCCAGAA | KX669268.1 |
| AAAAAACCAGCCCA | KX669268.1 |
| ACACAACGAGGGAA | KX669268.1 |
| CACTCACGCATTCT | KX669268.1 |
| ACCATCAACCCCAA | KX669268.1 |
| AGGGCTGTGATGA | KX669268.1 |
| GAAGCCGATATCCC | KX669268.1 |
| GTATGAGGAAGGGG | KX669268.1 |
| GGCGTAGGAAACAT | KX669268.1 |
| TTCGGGGTGCTT | KX669268.1 |
| AGGAAGCGAGAAGA | KX669268.1 |
| GGTTTGGTTGAGTG | KX669268.1 |
| CACCCCAACCGAAA | KX669268.1 |
| TTCGGGGTGCTT | KX669268.1 |
| TGGGGTGTCCTCGATTTTAG | KX669268.1 |
| CATGGTAATGCCTGCTGCTA | KX669268.1 |
| ATCACCTGAGCCCACCATAG | KX669268.1 |
| GGAACCCTGTTGCTACGAAA | KX669268.1 |
| CCATAGAAGCCTCCACCAAA | KX669268.1 |
| CCAAGTTTTATGGCGAGAGC | KX669268.1 |
| AACAGTGACACCCACTCTTCC | NC_009144.3 |
| TTACTCCTTGGAGGCCATGT | NC_009144.3 |
| GTATAGGGAATGAAAAACTAAGCATGAG | NC_009150.3 |
| GGGCAGACGGGAAACTGA | NC_009150.3 |
| CCGAAGGCTGCAGCTTTG | NC_009156.3 |
| TCCACCTCCATCCTCTTCCA | NC_009156.3 |
| GCCCGTCACCCTCCTTAAAT | KX669268.1 |
| GCCGAGTTCCTTTTACTTCTTTT | KX669268.1 |
| ATCCTAATGGTGCAACCGCT | KX669268.1 |
| TCCGCTTATTAGGAGGACTGAGA | KX669268.1 |
| TCCTAGCAGAATACGCAAACA | KX669268.1 |
| GTTGGGTTTGGTTGAGACCG | KX669268.1 |
| CCTACACTTCCACGACCACA | KX669268.1 |
| GGCGATTGTTGATTAGTCGGT | KX669268.1 |
| TATTCGCCTCTTTCGCTACCC | KX669268.1 |
| AAGGCTCAGAAGAAGCCAGAG | KX669268.1 |
| TGACTCCCCTACTACTCCTGTC | KX669268.1 |
| TGGAGTAGGGCTGAGACTGGT | KX669268.1 |
| ACTAGAACACTCCACCAGCG | KX669268.1 |

| | |
|---|---|
| ATGGGGGTTTAGTGCTGATTGT | KX669268.1 |
| TATTCTCCCCCGACCTCCTA | KX669268.1 |
| CGAGGATTGGGACACGTAGTT | KX669268.1 |
| ATCCAAACGTGGGGGTTTCT | KX669268.1 |
| TCTAGGGGGATGCCTGTCTA | KX669268.1 |

Note

unused in S2
unused in S2

unused in S2
